# Tubulin polyglutamylation modulates Golgi morphodynamics and neurite branching during neuronal morphogenesis

**DOI:** 10.64898/2026.04.13.718193

**Authors:** Chih-Hsuan Hsu, Alexander Josiah Kinrade, Sarah Cohen

## Abstract

Neurons establish functional networks through morphological remodeling during neuronal differentiation. Microtubule polyglutamylation is a key microtubule post-translational modification that is highly enriched during this process and plays an important role in differentiation. However, how remodeling of organelle features such as morphology, distribution and interactions depend on tubulin polyglutamylation during neuronal differentiation remain unclear. Here, we employed multispectral imaging combined with quantitative 3D organelle analysis to comprehensively profile eight organelles simultaneously in human induced pluripotent stem cell-derived neurons. We discovered that depletion of tubulin polyglutamylation induces pronounced alterations in somatic Golgi morphology and associated organelle interactions. In addition, Golgi-derived compartments in proximal neurites exhibited altered morphology and dynamics, namely decreased retrograde directionality. These changes were accompanied by increased neurite branching and tortuosity. Together, our findings reveal a previously unrecognized role for tubulin polyglutamylation in coordinating organelle organization with neurite architecture, providing a mechanistic link between tubulin post-translational modification, Golgi morphology, dynamics, and neuronal morphogenesis.

## INTRODUCTION

During neuronal differentiation, neurons undergo a series of morphological changes to establish highly polarized cellular architecture, including axons and dendrites, to form functional neural networks (Ho and Gupton, 2021; Menon and Gupton, 2018). These morphological changes require coordinated remodeling of the cytoskeleton, plasma membrane, and intracellular organelles (Hanus and Ehlers, 2016; Koppers and Farías, 2021). As a major cytoskeletal component, microtubules (MTs) play crucial roles in this process by providing structural support, serving as tracks for intracellular transport, and governing organelle organization and membrane trafficking (Burute and Kapitein, 2019; Schmoranzer and Simon, 2003). Microtubule dynamics and functions are regulated by diverse tubulin isotypes and post-translational modifications (PTMs), collectively known as the “tubulin code” (Janke and Magiera, 2020; Roll-Mecak, 2020), which has been shown to play important roles in neuronal development (Hsu et al., 2026; Park and Roll-Mecak, 2018). Tubulin polyglutamylation is one of the key tubulin PTMs that markedly accumulate in neurons during differentiation (Audebert et al., 1994; Hsu et al., 2026). It can function as a signal to direct organelle positioning, particularly of the endoplasmic reticulum (ER), and thereby potentially influence the distribution of other organelles through membrane contact sites (Zheng et al., 2022).

Studies have shown that metabolic reprogramming accompanies neuronal differentiation, a process often associated with changes in organelle organization and function, especially for mitochondria (Ordureau et al., 2021; Zanellati et al., 2026; Zhang et al., 2022; Zheng et al., 2016). In parallel, extensive multi-organelle rearrangement occurs across selective autophagy and secretory trafficking, involving coordinated changes among ER, Golgi, lysosomes and mitochondria (Hoyer et al., 2024; Ordureau et al., 2021; Wang et al., 2023). Our recent studies have shown that tubulin PTMs such as tubulin acetylation coordinate organelle organization and are important for maintaining organelle homeostasis during neuronal differentiation (Hsu et al., 2026). Although tubulin polyglutamylation and organelle remodeling have both been implicated in neuronal differentiation, no systematic analysis has been performed to understand how tubulin polyglutamylation contributes to the global landscape of multi-organelle remodeling to support the development of complex neuron architecture.

Multispectral imaging (MSI) has increasingly been applied as a systems-level approach for multi-organelle analysis, enabling comprehensive characterization of organelle features at the single-cell level (Arribat et al., 2020; Hsu et al., 2026; Rhoads et al., 2025; Valm et al., 2017; Zanellati et al., 2026; Zheng et al., 2022; Zimmermann et al., 2024). This approach overcomes the spectral channel limitations of conventional fluorescence imaging while avoiding the low throughput, labor-intensive preparation, and harsh fixation required for electron microscopy, and retains the speed and sensitivity of confocal microscopy (Zanellati et al., 2024). Here, we applied MSI together with an in-house computational analysis pipeline “infer-subc” to examine how eight organelles remodel in response to depletion of tubulin polyglutamylation in human induced pluripotent stem cell (hiPSC)-derived neurons (iNeurons). We discovered that tubulin polyglutamylation predominantly affects the somatic Golgi among the organelles examined, accompanied by altered organelle interactions involving the Golgi. In addition, tubulin polyglutamylation depletion changes the dynamics of Golgi-derived compartments in neurites. We further find that these alterations are associated with increased neurite branching. Together, these findings reveal a previously uncharacterized mechanism linking tubulin polyglutamylation-dependent regulation of organelle organization to neurite morphology.

## RESULTS AND DISCUSSION

### Multispectral imaging reveals organelle remodeling in response to tubulin polyglutamylation during neuronal differentiation

Our recent work demonstrated that coordinated remodeling of multiple organelles is a hallmark of neuronal differentiation (Zanellati et al., 2026). To study organelle remodeling during neuronal differentiation, we used a hiPSC NGN2 differentiation system to generate cortical neurons (iNeurons) in a controlled and rapid manner (Fig.1A). We previously found that during neuronal differentiation, tubulin polyglutamylation levels become highly enriched and show preferential subcellular accumulation in proximal neurites compared with the soma (Hsu et al., 2026; Fig. 1B). Tubulin polyglutamylation is catalyzed by Tubulin Tyrosine Ligase Like (TTLL) enzymes that add glutamate residues to the C-terminal tails of tubulin. TTLL1 serves as a primary α-tubulin polyglutamylase responsible for extending glutamate side chains on microtubules and is highly expressed in the brain (Janke et al., 2005; van Dijk et al., 2007). To investigate how tubulin polyglutamylation regulates organelle networks during neuronal differentiation, we reduced polyglutamylated tubulin levels by lentiviral shRNA-mediated knockdown of TTLL1. shRNA was transduced immediately after NGN2 induction (day 0), and cells were collected at day 7, achieving efficient knockdown as verified by Western blotting (Fig. 1C) and immunofluorescence staining (Fig. 1D).

**Fig. 1.**
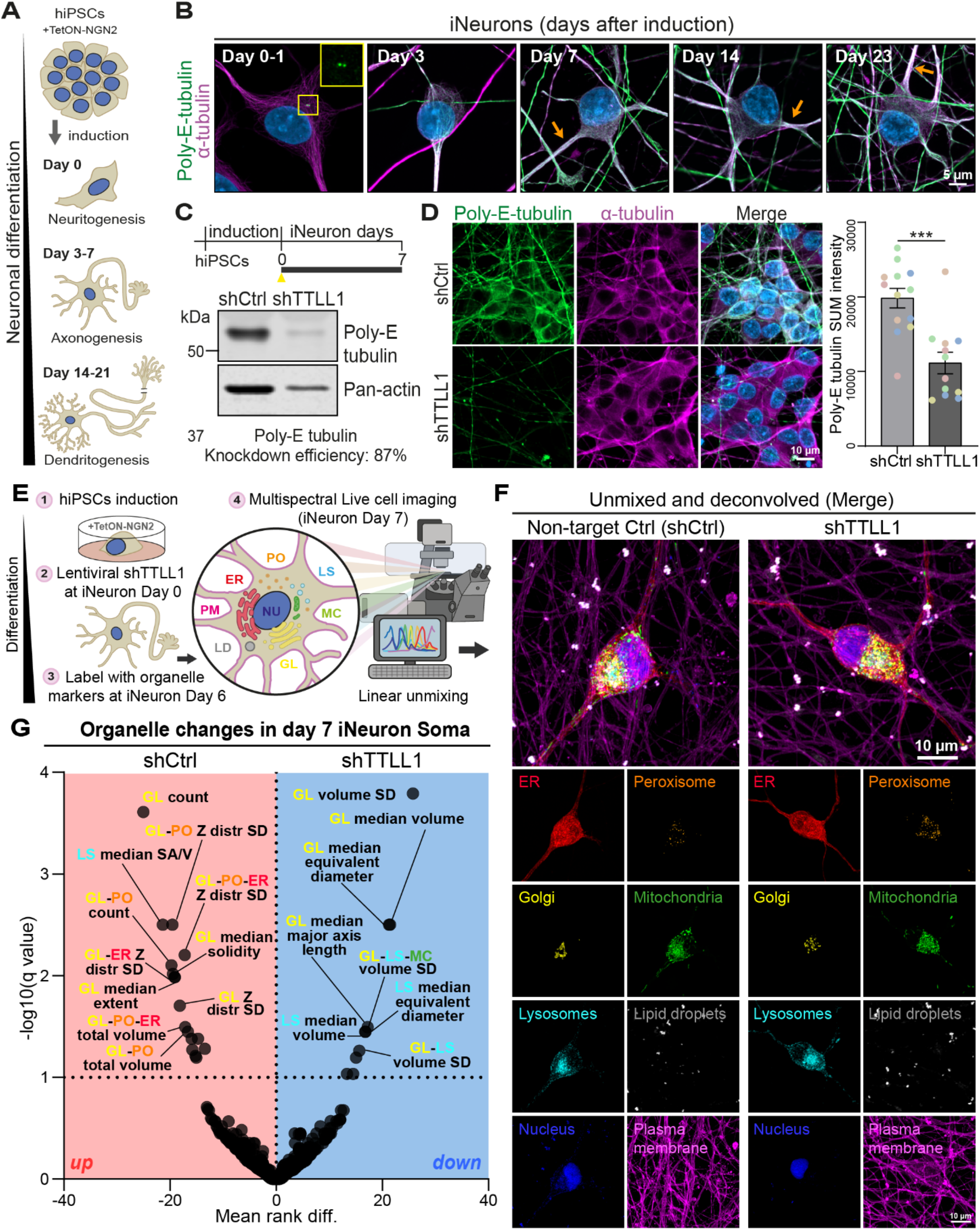
Multispectral imaging reveals organelle remodeling in response to tubulin polyglutamylation during neuronal differentiation. (A) Schematic, adapted from (Hsu et al., 2026), illustrating the NGN2-induced neuronal differentiation process across distinct morphogenesis stages. Human TetON-hNGN2-KOLF2.1J iPSCs were differentiated into cortical neurons (iNeurons) driven by NGN2 induction. (B) Airyscan Z-stack confocal images with maximum intensity projections of methanol fixed iNeurons at day 0-1, 3, 7, 14, and 23 post induction. Cells were immunolabeled for polyglutamylated tubulin (green) and α-tubulin (magenta). Nuclei were labeled with DAPI (blue). (C) Schematic diagram of non-target shRNA (shCtrl) versus TTLL1 (shTTLL1) lentivirus transduction and cell lysate collection timelines, where yellow arrowheads mark the day of viral transduction and bars denote the duration from transduction to sample collection. Western blot analysis shows correlating polyglutamylated tubulin protein levels. (D) Representative immunofluorescence confocal images show shCtrl-versus shTTLL1-treated day7 iNeurons fixed and stained for polyglutamylated tubulin (green), α-tubulin (magenta) and DNA (DAPI, blue). Bar graph shows the quantification of tubulin polyglutamylation SUM projected signal intensity from five biological replicates (color-coded), demonstrating decreased tubulin polyglutamylation in shTTLL1-compared with shCtrl-treated iNeurons. Unpaired two-tailed t test. Error bars represent ±SEM. ***, p<0.001. (E) Schematic of multispectral imaging workflow, adapted from (Hsu et al., 2026). (F) Individual channels from processed 3D multispectral images display eight organelles (endoplasmic reticulum (ER), peroxisomes (PO), Golgi (GL), mitochondria (MC), lysosomes (LS), lipid droplets (LD), nucleus (NU), and plasma membrane (PM)) separately. (G) Volcano plot illustrating soma-specific differences in 3D organelle organization (383 metrics) between shCtrl- and shTTLL1-treated day7 iNeurons. Statistical significance was determined using a 10% false discovery rate (FDR) threshold. SD, standard deviation. distr, distribution. n (cells)=39 shCtrl and 46 shTTLL1 from three biological replicates. Asterisks denote q values, * : q < 0.1,** : q < 0.05,*** : q < 0.01,**** : q < 0.001.

Next, we implemented a systematic multispectral microscopy technique to comprehensively capture 3D organelle organization under conditions of tubulin polyglutamylation perturbation. iNeurons treated with non-targeting (shCtrl) versus TTLL1-targeting (shTTLL1) shRNA were labeled with markers for eight organelles – endoplasmic reticulum (ER), peroxisomes (PO), Golgi (GL), mitochondria (MC), lysosomes (LS), plasma membrane (PM), lipid droplet (LD), and nucleus (NU) – and imaged at day 7 to visualize organelle organization (Fig. 1E and 1F). Single timepoint Z-stacks were acquired, linearly unmixed and deconvolved, followed by instance segmentation of organelles in each channel (Fig. S1A and S1B). We analyzed organelle features related to morphology, inter-organelle communication, and spatial distribution using the infer-subc pipeline, generating 5439 metrics, from which a curated subset of 383 was selected for downstream analysis. We found 28 metrics that were significantly different between shCtrl- and shTTLL1-treated day 7 iNeurons at a false discovery rate (FDR) of 10%, of which more than 78% of identified metrics were associated with Golgi-related changes (Fig. 1G, Table S1). We identified that depleting tubulin polyglutamylation resulted in distinct organelle remodeling, primarily in the soma and largely associated with Golgi-related features.

### Tubulin polyglutamylation-dependent organelle remodeling predominantly involves somatic Golgi

The curated 3D organelle multispectral imaging analysis revealed that the most significant changes observed were in Golgi morphology in the soma of shCtrl-versus shTTLL1-treated day 7 iNeurons. We identified an increase in the number of Golgi structures, while the size of individual Golgi structures decreased, resulting in an overall unchanged total Golgi volume and reduced Golgi size variability (Fig. 2A and 2B). In parallel, the Golgi shape became more fragmented, less irregular, and with fewer elongations, as indicated by decreased major axis length and equivalent diameter, and increased extent and solidity (Fig. 2C). In addition to Golgi, lysosome morphology was also significantly altered, with changes such as decreased individual lysosome size, increased surface area-to-volume ratio, and reduced major axis length and equivalent diameter, indicating a shift toward smaller and more compact lysosomes (Fig. 2B and 2C). Notably, distinct tubulin post-translational modifications exhibit differential effects on organelle remodeling, especially in lysosome morphology upon tubulin modifying enzyme depletion. In contrast to depleting tubulin polyglutamylation, tubulin acetylation depletion results in enlarged, circular lysosomes (Hsu et al., 2026). As these organelle organization changes were identified using the same multispectral imaging approach, this comparison highlights both the specificity of individual tubulin modifications in shaping organelle architecture and the sensitivity of the method to resolve distinct subcellular phenotypes. Next, we quantified lateral (XY) and axial (Z) organelle distributions, representing the spread of organelles from the nucleus to cell border and along the vertical axis of the cell from the bottom to the top, respectively. The lateral distribution of the Golgi remained unchanged, whereas Golgi showed increased variability in axial distribution in shTTLL1-treated iNeurons (Fig. 2D).

**Fig. 2.**
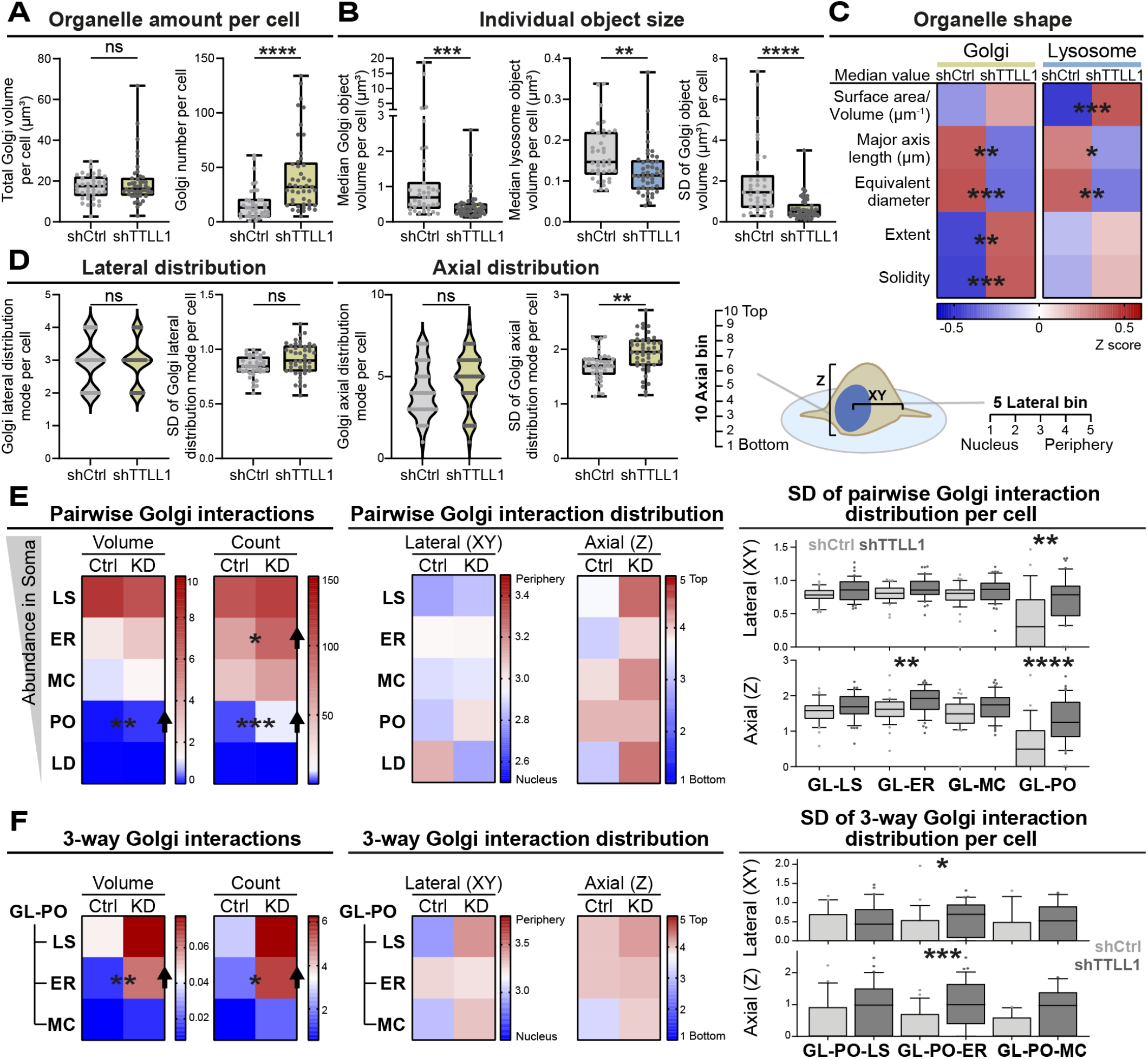
Tubulin polyglutamylation-dependent organelle remodeling occurs predominantly involved Golgi. (A-B) Box plots comparing shCtrl-versus shTTLL1-treated day 7 iNeurons for Golgi total volume (A, left), Golgi object number (A, right), individual Golgi object median size (B, left), individual lysosome median size (B, middle), and standard deviation (SD) values of Golgi object volumes (B, right). Data points represent single cells. Asterisks denoting q values from Fig. 1G. (C) Heatmaps displaying the organelle shape metrics for Golgi and lysosomes. Each metric is Z-score-normalized with asterisks denoting q values from Fig. 1G. (D) Schematic of organelle distribution measurements along the XY and Z axes (right), adapted from (Hsu et al., 2026). Quantification of the mode and standard deviation (SD) values of Golgi across each dimension (left). Asterisks denote q values from Fig. 1G. (E) Heatmaps of the total volume (μm^3^) and number of pairwise soma-specific Golgi interactions per cell in shCtrl-versus shTTLL1-treated day 7 iNeurons, ordered by organelle volume abundance (left). Pairwise Golgi interaction distribution mode along the lateral and axial are displayed in heatmaps (middle), with boxplots analysis of lateral or axial distribution standard deviation (SD) values for each Golgi pairwise interactions (right). Asterisks indicate q-values from Fig. 1G.(F) Three-way interactions involving Golgi and peroxisomes are quantified by total volume (μm^3^) and number (left), interaction distribution mode (middle) and the standard deviation (SD) of lateral and axial distributions (right) comparing between shCtrl- and shTTLL1-treated day 7 iNeurons. Asterisks indicate q-values from Fig. 1G.

Our recent work demonstrates that remodeling of inter-organelle contact networks rewires lipid metabolism to support neuronal differentiation (Zanellati et al., 2026). However, no studies have systematically characterized how tubulin polyglutamylation mediates these inter-organelle interactions during neuronal differentiation. To address this, we quantified organelle interactions through the overlapping regions between segmented organelle objects, referred to as organelle interaction sites or “contacts” as previously described (Hsu et al., 2026; Rhoads et al., 2025; Zanellati et al., 2026). Among all examined Golgi-organelle pairs, Golgi-lysosome interactions were the most abundant. However, Golgi-peroxisome interactions showed the largest changes upon depleting tubulin polyglutamylation, with increases in both the number and volume of interactions (Fig. 2E, left). The overall spatial distribution of interactions was not significantly altered (Fig. 2E, middle). Nevertheless, Golgi-peroxisome interactions became more randomly distributed in both the lateral and axial directions (Fig. 2E, right). One possible explanation is that Golgi fragmentation and dispersal increase opportunities for Golgi-peroxisome interactions. In addition, Golgi-peroxisome interplay has been implicated in supporting Golgi lipid composition and secretory function (König et al., 2025). Thus, altered Golgi architecture may increase reliance on Golgi-peroxisome interactions as a compensatory response to maintain Golgi lipid composition and support secretory demands. Analysis of three-way organelle contacts revealed the largest changes in interactions involving Golgi and peroxisomes. While Golgi-peroxisome-lysosome contacts were the most abundant, Golgi-peroxisome-ER contacts showed significant increases in both interaction number and volume in shTTLL1-treated iNeurons (Fig. 2F, left). Though Golgi-peroxisome-ER interactions remained unchanged spatially (Fig. 2F, middle), they exhibited increased spatial variability (Fig. 2F, right).

In summary, depletion of polyglutamylated tubulin resulted in prominent Golgi remodeling, characterized by increased Golgi fragmentation and greater variability in axial distribution. In addition to these morphological and spatial changes, Golgi-peroxisome interactions increased and became more randomly distributed. Moreover, interactions among Golgi, peroxisomes, and ER were also enhanced. As both peroxisomes and ER play central roles in lipid metabolism, increased Golgi interactions with these organelles may reflect enhanced coordination to meet lipid demands required for membrane trafficking.

### Reducing tubulin polyglutamylation alters trans-Golgi morphology and dynamics in neurites

In our multispectral imaging analysis, we observed increased fragmentation of the somatic Golgi in shTTLL1-treated iNeurons based on trans-Golgi labeling. Additionally, immunofluorescence staining of endogenous cis- and trans-Golgi markers using Airyscan imaging showed that both compartments appeared dispersed, suggesting a disruption of overall Golgi organization (Fig. S2A) that may impact the spatial organization of neuronal secretory trafficking. In addition to the somatic Golgi, which supports intracellular secretory pathways and primary dendrite specification, neurons contain Golgi-derived compartments within neurites. Golgi outposts in proximal neurites contribute to microtubule nucleation, neurite branching, and migration, while Golgi satellites are smaller, wider spread and serve as local processing and trafficking hubs (Nakagawa, 2024; Ori-McKenney et al., 2012). Polyglutamylated microtubules are highly enriched in proximal neurites compared with the soma, raising the possibility that tubulin polyglutamylation depletion may locally affect the dynamics of these Golgi-derived compartments in proximal neurites. Additionally, no significant changes were detected from the 122 neurite-specific curated subset (Fig. S2B, Table S1) across six organelles, supporting that the Golgi is the most affected. However, the strong somatic Golgi signal may limit detection of dim Golgi-derived structures in neurites in the multispectral imaging scheme.

To better examine neurite-specific Golgi, we expressed the trans-Golgi marker, pBac-mOxvenus-SIT (Sialyltransferase), alone and analyzed the dynamics of neurite-specific Golgi puncta (Fig. 3A, Movie 1). Here we identified that depletion of tubulin polyglutamylation biases Golgi-derived compartment trafficking toward anterograde movement with less retrograde trafficking compared to shCtrl-treated iNeurons (Fig. 3B). The decreased speed and increased turning angle indicate less directed and more irregular movement. While the median value of track duration remained unchanged, the mean track duration increased, suggesting an increased proportion of longer-lasting tracks in shTTLL1-treated neurites. This shift is accompanied by morphological alterations, including increased circularity and solidity (Fig. 3C), indicating a transition toward a more vesicular, transport-ready state. These changes may reflect a change in the composition of Golgi-derived compartments in neurites, potentially favoring smaller, satellite-like structures over more elongated outpost-like elements. Decreased variability in circularity and object area further suggests reduced morphological heterogeneity in shTTLL1-treated neurites. The altered dynamics of trans-Golgi compartments in shTTLL1-treated neurites suggest dysregulated Golgi organization not only at the soma but also within neurites, together with reduced inward vesicular delivery, pointing to an impact on secretory pathway including altered distribution of newly synthesized proteins and lipids.

**Fig. 3.**
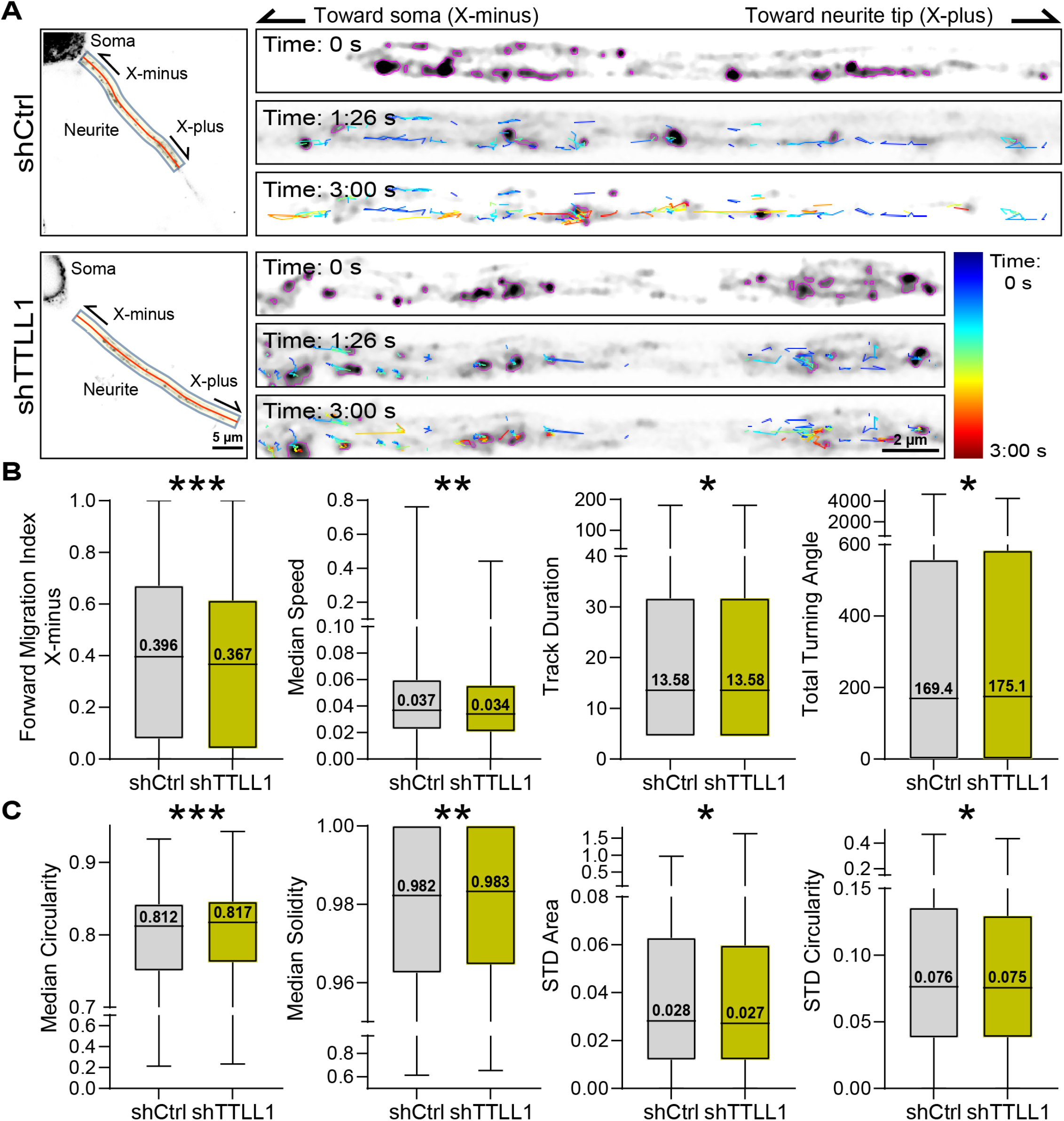
Reducing tubulin polyglutamylation alters trans-Golgi morphology and dynamics in neurites. (A) Day 7 shCtrl-versus shTTLL1-iNeurons with trans-Golgi labeling were imaged using Airyscan for 3 min at 4.53 s intervals. Neurite regions selected for analysis were tracked and straightened from the soma toward the distal neurite within the field of view, as indicated in the representative images. Tracks are color-coded by time. (B-C) Box plots of trans-Golgi dynamics, including forward migrating index X-minus, median speed, track duration, and total turning angle (B). For track duration, Mean ± SEM values are: 26.23 ± 0.5 shCtrl; 28.16± 0.8 shTTLL1. Morphological features of tracked objects, including circularity, solidity, area standard deviation (STD) are shown in (C). Median values are indicated, and values represent population-level summaries across multiple neurites; n (neurites)=63 shCtrl (34 cells) and 38 shTTLL1 (22 cells) from three biological replicates. Box plots depict the median, interquartile range, and range; asterisks denote p values determined by a Bonferroni-corrected statistical randomization test. *, p<0.05; **, p<0.01; ***, p<0.001; ****, p<0.0001.

### Loss of tubulin polyglutamylation alters neuronal morphology and increases neurite branching

Small Golgi outposts were previously found to selectively partition into longer, more complex dendrites to support directed secretory trafficking required for asymmetric dendrite growth and morphogenesis (Horton et al., 2005). In addition, α-tubulin polyglutamylation, regulated by doublecortin (DCX), has been implicated in restricting neurite branching during early neuronal development, as DCX knockout iNeurons exhibit increased neurite initiation without major changes in microtubule dynamics (Sébastien et al., 2025). The reduced retrograde dynamics of Golgi-derived compartments observed in neurites upon TTLL1 depletion suggested increased or less spatially restricted membrane delivery through secretory trafficking. Therefore, we next examined neuronal morphology and neurite network organization. iNeuron cultures are highly intertwined, making it challenging to distinguish neurites from individual cells. Cellmasks generated from the multispectral dataset were leveraged, as sparse labeling enabled segmentation and quantification of individual neurons (Fig. 4A, left). Tubulin polyglutamylation depletion did not alter soma volume but significantly increased the number of primary neurites directly extending from the soma (Fig. 4A, right).

**Fig. 4.**
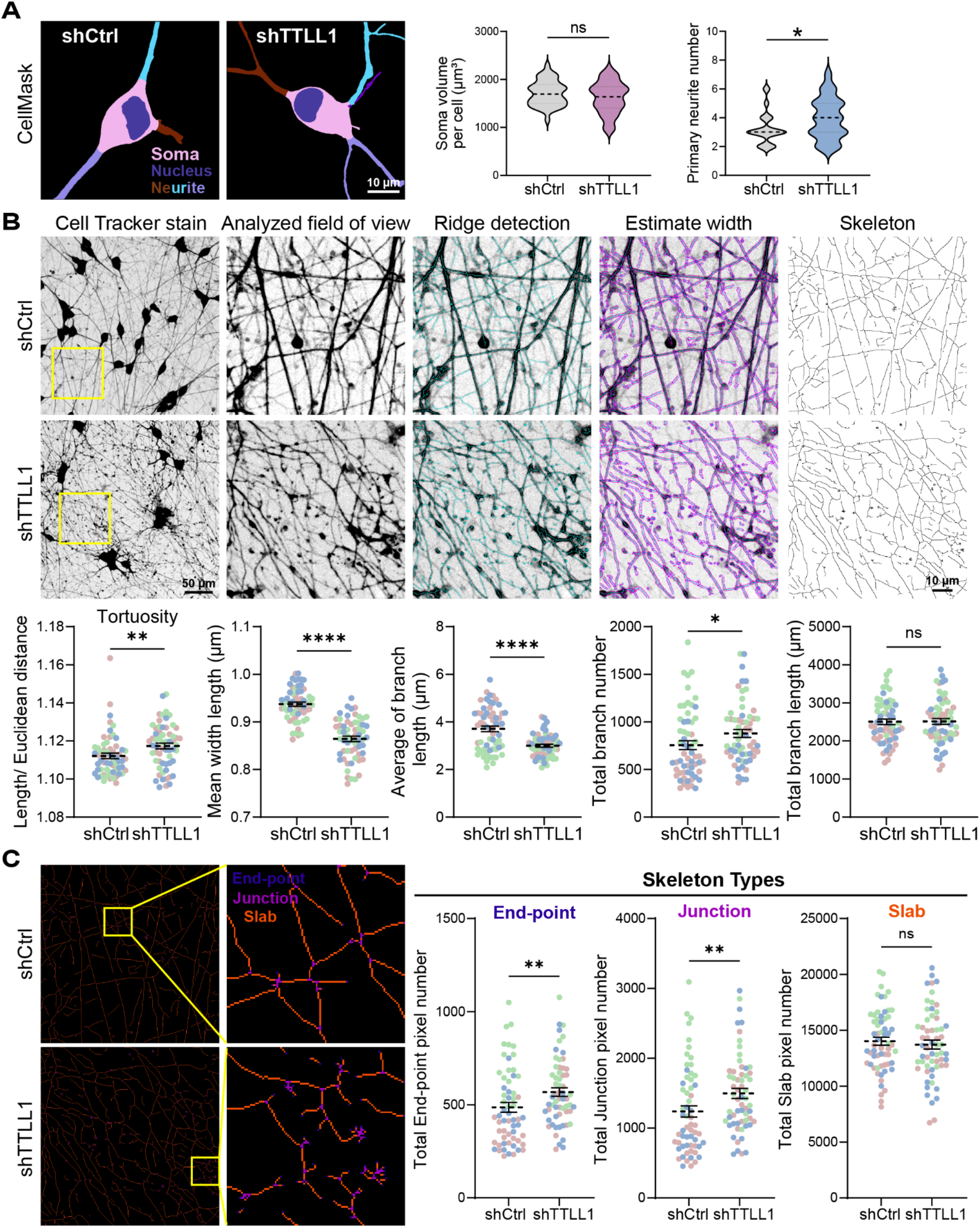
Loss of tubulin polyglutamylation alters neuronal morphology and increases neurite branching. (A) Masks derived from multispectral processing were used to quantify soma volumes and primary neurite number between shCtrl-versus shTTLL1-treated day 7 iNeurons. n (cells)=39 shCtrl and 46 shTTLL1 from three biological replicates. Statistical significance was determined using an unpaired t-test with *, p<0.05. (B) CellTracker-labeled day 7 shCtrl-versus shTTLL1 iNeurons were processed by ridge detection and skeletonization to quantify tortuosity index (ratio of skeletonized length to Euclidean distance), mean width length, average branch length, total branch number and total branch length, data represent mean ± SEM. n (analyzed field of view)=60 shCtrl and 60 shTTLL1 from three biological replicates, with data points color-coded by replicate. Statistical significance for mean width length and total branch length was determined using an unpaired t-test. For tortuosity index, average branch length, and total branch number, Mann-Whitney-test were applied. *, p<0.05; **, p<0.01; ***, p<0.001; ****, p<0.0001. In the representative images, the yellow box indicates the cropped region for analysis, cyan lines mark detected neurite ridges, and magenta outlines estimated neurite width. (C) Skeletonized images from (B) were analyzed for pixel counts of end-points, junctions, and slabs (right). Mann–Whitney test was used for statistical analysis. *, p<0.05; **, p<0.01; ***, p<0.001; ****, p<0.0001.

Next, to examine the effect of tubulin polyglutamylation on neurite network organization, iNeuron cultures were labeled with CellTracker and analyzed by skeletonization. Depleting polyglutamylated tubulin increased neurite tortuosity with decreased neurite width. Comparable total neurite content per field of view was analyzed, but tubulin polyglutamylation depletion resulted in a greater number of shorter and more fragmented branches (Fig. 4B). To further characterize neurite networks, skeleton features, including endpoints, slabs, and junctions, were analyzed. While the number of slabs remained unchanged, polyglutamylated tubulin depletion resulted in an increased number of endpoints and junctions, indicating enhanced branching complexity (Fig. 4C).

In conclusion, multispectral imaging revealed that tubulin polyglutamylation mediated by TTLL1 regulates the morphology, distribution, and interactions of the somatic Golgi among six major organelles, as well as the morphology and dynamics of Golgi-derived compartments in proximal neurites. Together with its elevated enrichment in proximal neurites, these findings support a spatially specific role for polyglutamylated microtubules in controlling Golgi-derived compartment dynamics in neurites. The neuronal network phenotype we observed, characterized by increased branching and reduced straightness, may be explained by intracellular dysregulation linking tubulin polyglutamylation-dependent control of Golgi organization to neurite architecture.

## MATERIALS AND METHODS

### Culture of human iPSC derived neurons

Human KOLF2.1J wildtype (wt) induced pluripotent stem cells (hiPSCs) were obtained from Bill Skarnes (The Jackson Laboratory) and were used to generate a pBac-TetON-hNGN2 iPSCs stable cell line as previously described (Fernandopulle et al., 2018; Zanellati et al., 2026). Briefly, KOLF2.1J wt hiPSCs were transfected with 500 ng piggyBac plasmid encoding a bipartite TetON-hNGN2 cassette for a well of a 6-well plate (plasmids gifted from Michael Ward; NINDS). Following transfection, cells were selected for stable genomic integration using puromycin (0.7 µg/ml). Cells were maintained and cultured as adherent monolayers at 37°C in a humidified incubator with 5% CO_2_ under feeder-free conditions in StemFlex medium (A3349401; Gibco) on vitronectin (VTN-N)-coated 6-well plates (VTN-N Recombinant Human Protein, Truncated, A14700; Gibco). Plates were coated with vitronectin by diluting 10 µL vitronectin in 1 mL of 1x Dulbecco’s Phosphate-Buffered Saline (DPBS, 14190144; Gibco) per well and incubating for 2 hrs at 37°C. The hiPSCs were media changed every 1-2 days and were passaged upon reaching ∼80% confluency using ReLeSR (#100-0484; STEMCELL Technologies) according to manufacturer’s instructions in the presence of 0.5x ROCK inhibitor (RevitaCell Supplement, A2644501; Gibco).

### Differentiation into iNeurons

An established protocol was adapted to differentiate pBac-TetON-hNGN2 iPSCs into induced neurons (iNeurons)(Fernandopulle et al., 2018; Pantazis et al., 2022), with modifications (Hsu et al., 2026; Zanellati et al., 2026). Cultured hNGN2-iPSCs were dissociated into single cells using Accutase (A6964-100ML; Sigma-Aldrich) and cells (1.0-1.2×10^6^ per well of 6-well plates) were seeded in induction medium (IM) consisting of KnockOut™ DMEM/F-12 (12660012; Gibco) supplemented with 1x N-2 (17502048; Gibco), 1x MEM Non-Essential Amino Acids Solution (MEM-NEAA, 11140050; Gibco), 1x GlutaMAX (35050061; Gibco), 1x ROCK inhibitor (RevitaCell Supplement, A2644501; Gibco) and 0.5 μg/ml Doxycycline hyclate (D9891; Sigma-Aldrich). Twenty-four hrs after plating, media was replaced with IM (1x N-2, 1x MEM-NEAA, 1x GlutaMAX) with 1 μg/ml Doxycycline hyclate. After 48 hrs, 4×10^4^ induced cells were replated using Accutase onto each well of a 8-well Nunc Lab-Tek II chamber with 0.1 mg/mL poly-L-ornithine hydrobromide (Sigma; #P3655) in Borate buffer (10 mM Boric acid (Sigma B6768), 2.5 mM Sodium tetraborate (Sigma 221732), 7.5 mM Sodium chloride (Sigma S7653), 0.1 M Sodium hydroxide (Sigma 71463)) and 10 μg/mL Laminin (Gibco™ Laminin Mouse Protein, Natural, 1mg; 23-017-015) coated in IM (1x N-2, 1x MEM-NEAA, 1x GlutaMAX) with 0.5 μg/ml Doxycycline hyclate and 0.1x ROCK inhibitor (RevitaCell Supplement). 72 hrs after Doxycycline induction (defined as day 0 iNeurons), cultures were switched to cortical neuron culture media consisting of BrainPhys (05791; STEMCELL Technologies), 10 μg/mL BDNF (450-02; PeproTech), 10 μg/mL NT-3 (450-03; PeproTech), 2% B-27 (A1486701; Gibco), 1 μg/mL Laminin (23-017-015; Gibco) and supplemented with 0.5 μg/ml Doxycycline hyclate. iNeurons were cultured up to 21 days as adherent monolayers at 37°C in a humidified incubator with 5% CO_2_ with half medium changes every 2-3 days in doxycycline-free cortical neuron culture medium.

### iNeuron transduction

Lentiviral shRNA expression plasmids were from Sigma: MISSION® TRC-puro Non-Mammalian shRNA for control, TRCN0000048527 for TTLL1. Lentivirus was concentrated with Lenti-X Concentrator (Takara Bio 631231) in miliQ and cells were treated with virus diluted 1:2000 in cortical media. 24 hours after the transduction, medium was replaced with fresh cortical neuron culture media.

### Western blotting

Cells were lysed on ice in RIPA buffer (50 mM Tris, pH 8, 150 mM NaCl, 0.5% deoxycholate, 0.1% SDS, 1% NP-40 supplemented with 15 mM sodium pyrophosphate, 50 mM sodium fluoride, 40 mM β-glycophosphate, 1 mM sodium vanadate, 100 mM phenylmethylsulfonylfluoride, 1 μg/mL leupeptin, and 5 μg/mL aprotinin) supplemented with 1:100 Protease Inhibitor Cocktail (Sigma; P8340-5mL), 1:100 Phosphatase Inhibitor Cocktail 2 (Sigma; P5726), 1:100 Phosphatase Inhibitor Cocktail 3 (Sigma; P0044). Adherent samples were scraped, collected, and centrifuged at 13,000 xg for 5 min at 4°C to isolate the post-nuclear supernatant. Concentration of supernatant was measured by Pierce BCA Protein Assay Kits (Thermo Scientific; #23225), and the supernatant were then boiled with 6x Laemmli buffer containing β-mercaptoethanol and denatured for 10 min at 95°C. Equal protein amounts were loaded into each well of a 10% SDS-polyacrylamide gel and transferred onto 0.2 μm nitrocellulose membrane (Bio-Rad 162-0112) by wet transfer 1 h at 100 V for immunoblotting. Membranes were blocked in 5% milk in Tris-buffered saline (TBS) for 30 min at room temperature and then incubated with primary antibodies diluted in 5% milk in TBST overnight at 4°C. The next day, membranes were washed three times 5 mins with TBST, incubated with secondary antibodies diluted in 5% milk in TBST 1 h at room temperature, washed three times 5 mins with TBST, and imaged using an Odyssey CLx (LI-COR Biosciences). Western blots were quantified using LI-COR Image Studio. Background was subtracted for each quantified band. Knockdown efficiency was calculated by normalizing the signal intensity of polyglutamylated tubulin to pan-actin and then comparing the normalized values between knockdown and control groups.

### Antibodies

Antibodies used were: rabbit polyclonal anti-Polyglutamate chain (polyE), pAb (IN105)(AdipoGen Life Sciences AG-25B-0030-C050; 1:300 IF, 1:5000 WB), mouse against α-tubulin DM1A (Abcam #ab7291; 1:1000 IF, 1:10000 WB), mouse monoclonal against actin (SPM161) (Santa Cruz sc-56459; 1:5000 WB), chicken anti-MAP2 Polyclonal (Invitrogen PA1-10005; 1:1000 IF), mouse anti-GM130 (BD Biosciences 610822; 1:400 IF), and rabbit polyclonal Golgin 97 antibody (Abcam #ab84340;1:400 IF).

### Immunofluorescence staining and imaging of polyglutamylated microtubules

Cells were fixed with ice-cold 100% methanol for 10 mins at -20°C, then washed with DPBS 3 times. Cells were blocked using a blocking solution of 4% bovine serum albumin (BSA) in DPBS for 1 hour. Primary antibodies against polyglutamylated tubulin (rabbit, 1:300) and α-tubulin DM1A (mouse, 1:1000) were diluted in blocking solution and incubated with cells for overnight at 4°C. After primary antibody incubation, cells were washed with DPBS in three intervals of 5 minutes. Secondary antibodies, anti-rabbit 488 (1:1000) and anti-mouse 568 (1:1000), were diluted in blocking solution at room temperature for 1 hour while covered from light. After incubation, the secondary antibodies were washed out using three 5-minute DPBS washes and stained with DAPI 1:1000 for 10 mins. Labeled Cells were image using a Zeiss laser scanning microscope (LSM) 800 with a 63x 1.4 NA oil objective.

### Multispectral imaging of organelles

Day 6 iNeurons were co-transfected with six piggyBac organelle reporter plasmids together with a piggyBac transposase plasmid using Lipofectamine 2000 (11668027; Thermo Fisher Scientific) following a modified version of the manufacturer’s protocol. For each well of an 8-well chamber, plasmid DNA (120ng EIF1α-Transposase, 120 ng LAMP1::mTurquoise for lysosome, 120 ng Cox8::eGFP for mitochondria, 120 ng mApple::Sec61B for ER, 60ng SiT::oxVenus for Golgi, 25ng mOrange:: SKL for peroxisome, and 60 ng mTagBFP2::NLS for nucleus) was diluted in 25 µL Neurobasal media (12348017; Gibco). In parallel, 1 µL Lipofectamine 2000 was diluted in 25 µL neurobasal media and incubated for 5 min at room temperature. The diluted DNA and Lipofectamine 2000 solutions were then combined, properly mixed and incubated for 15 min at room temperature. Prior to transfection, cultures were washed and equilibrated with neurobasal medium. After a 45-min incubation with the transfection mixture at 37°C and 5% CO_2_, cultures were washed once and medium replaced with BrainPhys Imaging Optimized Medium (05796; STEMCELL Technologies) supplemented with 2% NeuroCult SM1 (05711; STEMCELL Technologies), 10 μg/mL BDNF (450-02; PeproTech), 10 μg/mL NT-3 (450-03; PeproTech), and 1 μg/mL Laminin (23-017-015; Gibco). On day 7 of iNeuron differentiation, 15 min prior to live-cell imaging, 1:2000 CellMask Deep Red (C10046; Invitrogen) was added to label the plasma membrane and 0.5 µM Lipi-Blue (LD01; DOJINDO Laboratories) was added to label lipid droplets. For single-label controls, each plasmid or dye was applied individually using the same DNA-to-Lipofectamine ratios or dye concentrations as in multi-labeled conditions.

### Multispectral imaging

Multispectral imaging was carried out as previously described (Hsu et al., 2026; Rhoads et al., 2025; Zanellati et al., 2026). Images were acquired using a Zeiss LSM 880 laser scanning confocal microscope on an inverted Axio Observer Z1 base equipped with a 34-channel GaAsP spectral detector and a live-cell incubation chamber (Carl Zeiss Microscopy). 405 nm (0.5%), 458 nm (3%), 514 nm (0.4%), 561 nm (0.3%), and 633 nm (0.2%) lasers were used simultaneously to excite cells labeled with one or multiple organelle markers. Emitted fluorescence was collected in lambda mode using a linear array of 34 photomultiplier tubes (PMTs) with 9 nm spectral bins spanning 410-695 nm, resulting in 32 intensity channels per image. Z-stack images were acquired by a plan-Apo 63x/1.4 NA oil-immersion objective with an XYZ voxel size of 0.07 µm × 0.07 µm × 0.39 µm. Z-stacks were acquired at a single time point with a pixel density of 796 × 796 per slice, and each Z-slice was captured in 1.97s. The number of Z-slices per image was determined on a cell-by-cell basis to capture the entire cell volume from basal to apical surfaces. All imaging was performed live and maintained at 37°C and 5% CO_2_ during imaging.

### Multispectral image processing

Using the linear unmixing algorithm from ZEN Black software version 2.3 (Carl Zeiss Microscopy), spectral Z-stack images were first processed into 8-channel images. Control images labeled with a single fluorophore corresponding to one of the eight organelle markers were acquired as described above and used to generate reference spectra. For each fluorophore, a representative region of fluorescence intensity was selected to define its reference spectrum. All reference spectra were then used as inputs and applied to all images across the full biological replicate dataset for linear unmixing, resulting in an eight-channel image in which each channel represents fluorescence intensity from a single fluorophore.

Linear unmixed images were subsequently deconvolved using Huygens Essential software version 23.04 (Scientific Volume Imaging, The Netherlands, http://svi.nl). Excitation and emission maxima (Ex/Em max) for each fluorophore were estimated from the lasers used during imaging and the corresponding reference spectra and applied for batch deconvolution of eight intensity channels using the Workflow Processor module.

For lateral (XY) distribution, segmented Z-stacks were summed across Z planes and divided into five equal bins with region 1 as the nucleus and regions 2-5 spanning from the nuclear edge to the cell periphery. For axial (Z) distribution, summed projections across XY axes were divided into ten equal bins from bottom to top (1 to 10) to quantify organelle volumes within each Z-section.

### Immunofluorescence staining of cis- and trans-Golgi compartments

Cells were fixed with ice-cold 100% methanol for 10 mins at -20°C, then washed with DPBS 3 times. Cells were blocked using a blocking solution of 4% bovine serum albumin (BSA) in DPBS for 1 hour. Primary antibodies against MAP2 (chicken, 1:1000), GM130 (mouse, 1:400) and Golgin97 (rabbit, 1:400) were diluted in blocking solution and incubated with cells overnight at 4°C. After primary antibody incubation, cells were washed with DPBS in three intervals of 5 minutes. Secondary antibodies anti-chicken 488 (1:500), anti-mouse 568 (1:500), and anti-rabbit 647 (1:500) were diluted in blocking solution and incubated at room temperature for 1 hour while covered from light. After incubation, the secondary antibodies were washed out with DPBS in three intervals of 5 minutes. DAPI was then added in DPBS (1:1000) and incubated at room temperature for 10 minutes. The DAPI solution was washed out and replaced with DPBS, and the samples were imaged using a Zeiss laser scanning microscope (LSM) 800 with a 63x 1.4 NA oil objective. Airyscan Z-stack or z-slice images were collected and processed using ZEN 2.3 Airyscan 3D processing software.

### Live-cell assay of trans-Golgi dynamics in neurites

Day 6 iNeurons were transfected with a single piggyBac organelle reporter plasmid with a piggyBac transposase plasmid using Lipofectamine 2000 (11668027; Thermo Fisher Scientific) following a modified version of the manufacturer’s protocol. For each well of a 4-well chamber, plasmid DNA (40 ng EIF1α-Transposase and 120 ng SiT::oxVenus) was diluted in 50 µL Neurobasal media. In parallel, 2 µL of Lipofectamine 2000 was diluted in 50 µL Neurobasal media. The diluted DNA and Lipofectamine 2000 solutions were then combined, properly mixed and incubated for 15 min at room temperature. Prior to transfection, cells were washed and equilibrated with neurobasal medium. After a 45 min incubation with the transfection mixture at 37°C and 5% CO_2_, cultures were washed once with Neurobasal media and replaced with cortical neuron culture media. Airyscan timelapse images were acquired on a Zeiss laser scanning microscope (LSM) 800 with a 63x 1.4 NA oil objective and processed using ZEN 2.3 Airyscan 2D Processing software. Timelapse images included a single Z-plane imaged every 4.53 seconds for 3 min (41 frames). Cells were imaged live and maintained at 37 C and 5% CO_2_ during imaging. The dynamics of trans-Golgi vesicles were quantified from 2D timelapse datasets using TrackMate and CellTracksColab (Ershov et al., 2022; Gómez-de-Mariscal et al., 2024). Timelapse images were processed prior to analysis. Briefly, neurite segments were isolated using the straighten function in FIJI and trans-Golgi vesicle signals were enhanced by subtracting background signal (radius = 25), employing a median filter (radius = 2) and using the enhance contrast function (saturate pixels = 0.35%) prior to vesicle segmentation. In TrackMate, the threshold detector (intensity threshold = 0.45) and LAP tracker (linking max distance = 3.5, radius penalty = 3, Y penalty = 15.0; gap closing max distance = 5.0, max frame gap = 2, radius penalty = 3.0, Y penalty = 15.0) were used. The CSV files from TrackMate were imported into CellTracksColab to visualize and analyze the tracking data. CellTracksColab v1.0.4 was used following the recommended data structure and analysis steps for the raw tracks. The metrics of trans-Golgi vesicle dynamics from CellTracksColab analysis include: Forward Migration Index X-Minus and Total Turning Angle. Forward Migration Index X-Minus is the fraction of the total path length resulting from movement in the negative x-direction. Total turning angle is the sum of the angles between each pair of consecutive direction vectors along the track duration.

### Neurite morphology assay

To visualize neuronal morphology, day 7 iNeurons were incubated with Cell Tracker Green (Invitrogen, C7025) (25 µM) for 15 min at 37°C and washed once with complete medium prior to imaging. Following dye incubation, Z-stacks of cells were collected using a Zeiss laser scanning microscope (LSM) 800 with a 20x objective. Z-stack images were maximum-intensity projected and then regions of interest were randomly selected, excluding soma-containing areas, and cropped to the field of view for analysis. Images were then processed using Enhance Local Contrast (CLAHE), followed by median filtering (radius = 2). Processed images were analyzed using the Ridge Detection plugin in Fiji to detect neurite ridges and estimate width (Steger, 1998; Wagner et al., 2017). Ridge-detected images were then converted to skeletonized images and analyzed using the Analyze Skeleton (2D/3D) plugin in Fiji (Arganda-Carreras et al., 2010; Doube et al., 2010; Polder et al., 2010).

### Statistics

The number of replicates and statistical tests are indicated in the figure legends. Each replicate was from a separate cell culture. For comparisons between two groups, T-tests or Mann-Whitney-test were used. For multiple-group comparisons, one-way ANOVA followed by appropriate post hoc tests (specified in the figure legends) was performed. For microscopy experiments, fields of view were selected at random, but image acquisition and analysis were not performed blinded to experimental conditions. For multispectral imaging datasets, a nonparametric Mann-Whitney U test and statistical significance was determined by a 10% false discovery rate. All statistical analyses, plots and heatmaps were conducted using GraphPad Prism 11 (RRID:SCR_002798) Windows, GraphPad Software, Boston, Massachusetts USA, www.graphpad.com. For trans-Golgi dynamic tracking, the statical significance was determined using the Bonferroni-corrected statistical outcomes from the CellTracksColab randomization test (Gómez-de-Mariscal et al., 2024).

## Acknowledgements

We thank Dr. Bill Skarnes (The Jackson Laboratory) for generously providing the KOLF2.1J cell line. We thank Dr. Michael Ward (NINDS) for kindly sharing piggyBac vectors. We thank Wendy Salmon (Hooker Imaging Core) for expert imaging advice. We thank Shannon Rhoads, Ngudiankama Rene Mfulama and Zachary Coman for assistance with the 3D organelle analysis pipeline (infer-subc). We thank Maria Clara Zanellati for helping with the neuron differentiation protocol.

## Footnotes

### Author contributions

C-H.H. and S.C. conceived the study, obtained funding, and wrote the paper. C-H.H. and A.J.K. performed the experiments and analyzed the data. S.C. supervised the project.

### Data and resource availability

The data generated in this study are available from the corresponding author upon reasonable request. Data is available at BioImage Archive under accession number S-BIAD3209.

## Supplementary information

**Figure S1.**
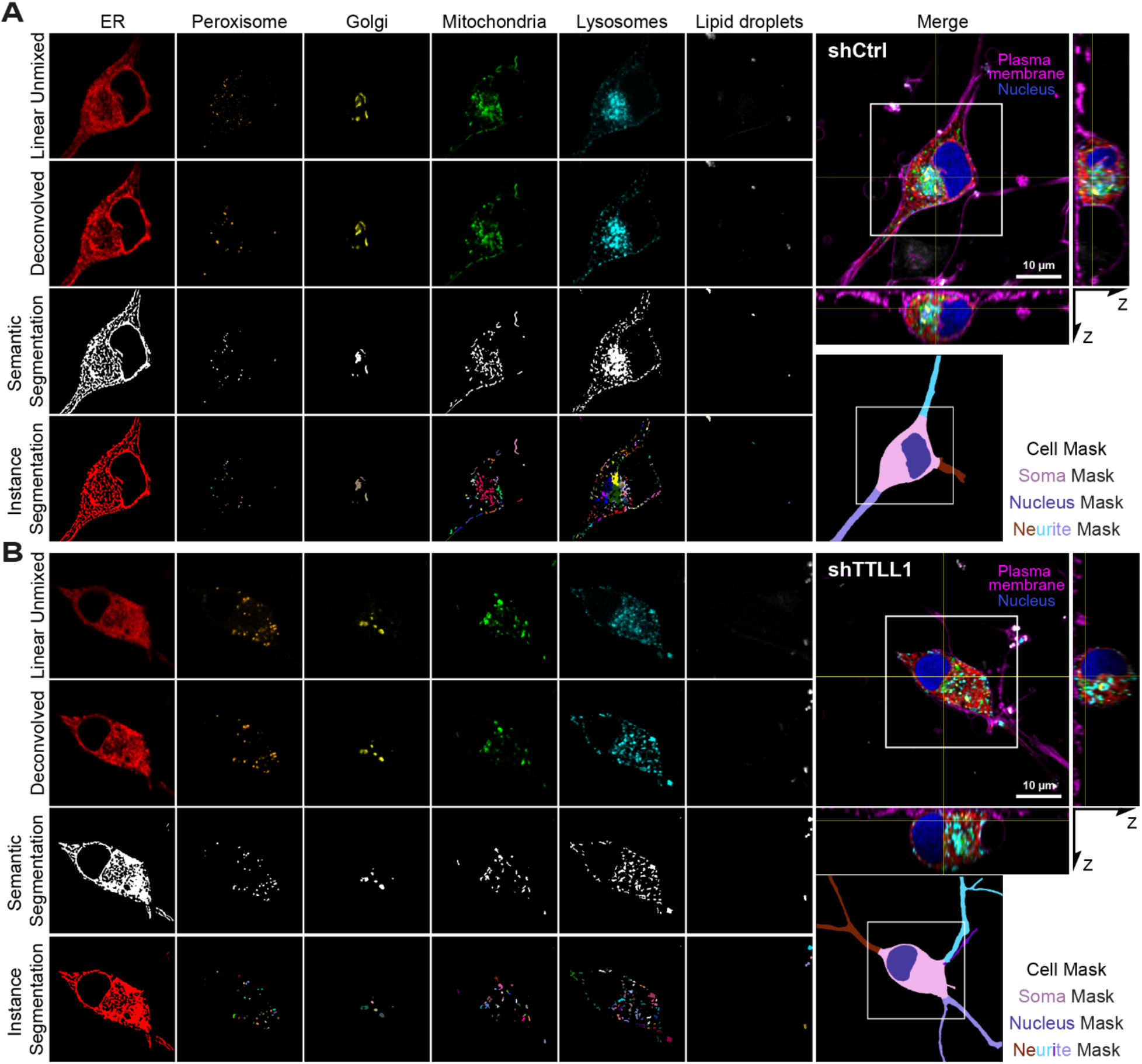
Representative images illustrating the multispectral processing steps: Linear unmixed fluorescence image after spectral separation, deconvolved images using Huygen deconvolution software, segmentation outputs and cell masks using infer-subc pipeline of day 7 iNeurons shCtrl (A) and shTTLL1 (B).

**Figure S2.**
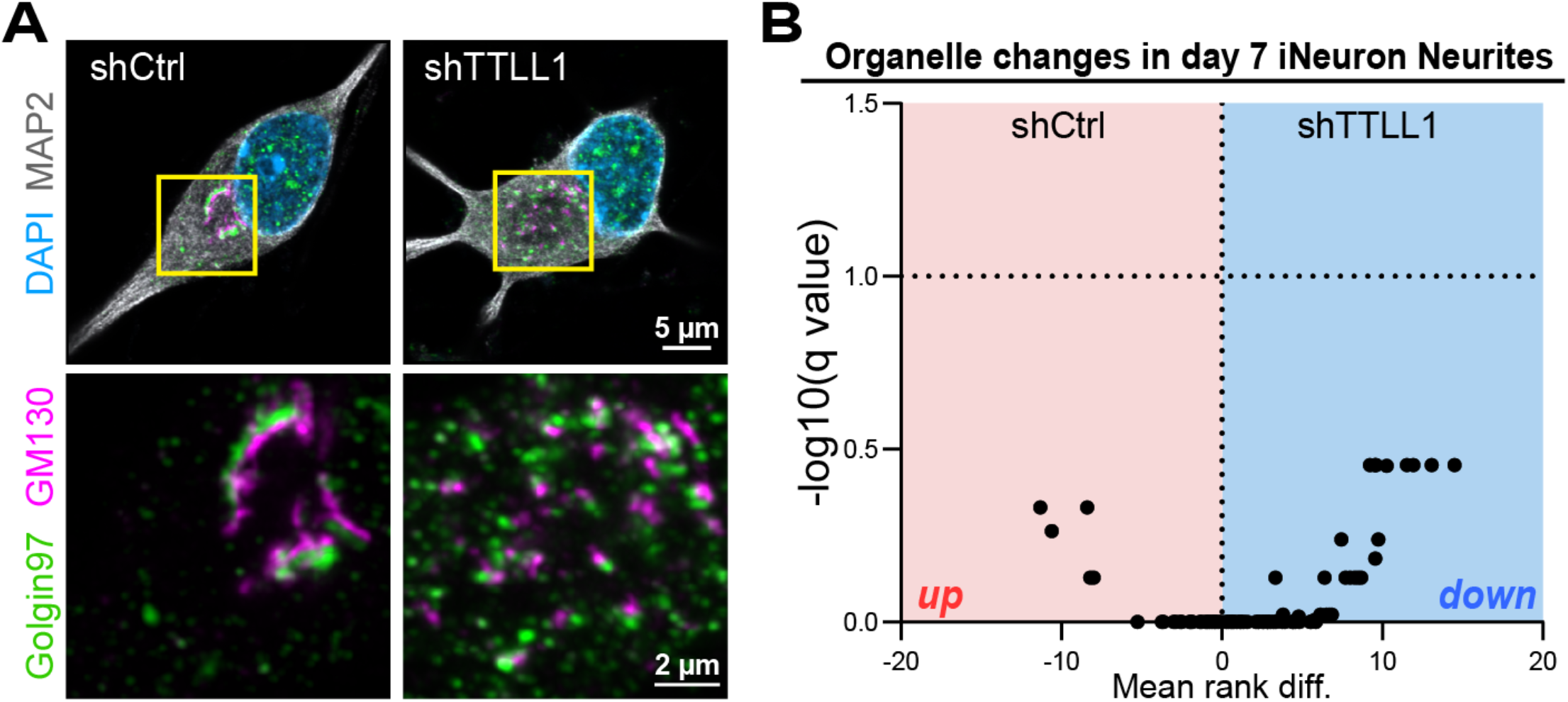
(A) Representative Airyscan images of shCtrl-versus shTTLL1-treated day 7 iNeurons stained for cis-Golgi (GM130), trans-Golgi (Golgin97), MAP2 and nuclei were labeled with DAPI. (B) Volcano plot illustrating neurite-specific differences in 3D organelle organization (122 metrics) between shCtrl- and shTTLL1-treated day7 iNeurons. Statistical significance was determined using a 10% false discovery rate (FDR) threshold. n (cells)=39 shCtrl and 46 shTTLL1 from three biological replicates.

